# Optogenetic relaxation of actomyosin contractility uncovers mechanistic roles of cortical tension during cytokinesis

**DOI:** 10.1101/2021.04.19.440549

**Authors:** Kei Yamamoto, Haruko Miura, Motohiko Ishida, Satoshi Sawai, Yohei Kondo, Kazuhiro Aoki

## Abstract

Actomyosin contractility generated cooperatively by nonmuscle myosin II and actin filaments plays essential roles in a wide range of biological processes, such as cell motility, cytokinesis, and tissue morphogenesis. However, it is still unknown how actomyosin contractility generates force and maintains cellular morphology. Here, we demonstrate an optogenetic method to induce relaxation of actomyosin contractility. The system, named OptoMYPT, combines a catalytic subunit of the type I phosphatase-binding domain of MYPT1 with an optogenetic dimerizer, so that it allows light-dependent recruitment of endogenous PP1c to the plasma membrane. Blue-light illumination was sufficient to induce dephosphorylation of myosin regulatory light chains and decrease in traction force at the subcellular level. The OptoMYPT system was further employed to understand the mechanics of actomyosin-based cortical tension and contractile ring tension during cytokinesis. We found that the relaxation of cortical tension at both poles by OptoMYPT accelerated the furrow ingression rate, revealing that the cortical tension substantially antagonizes constriction of the cleavage furrow. Based on these results, the OptoMYPT system will provide new opportunities to understand cellular and tissue mechanics.

## INTRODUCTION

Actomyosin contractility underlies force generation in a wide range of cellular and tissue morphogenesis in animals. Prominent examples include the tail retraction of directionally migrating fibroblasts and the constriction of a contractile ring during cytokinesis (Green, Paluch, and Oegema 2012; Ridley et al. 2003). The actin-rich cell cortex, a thin network underneath the plasma membrane, is also relevant to the actomyosin contractility involved in maintaining cell morphology; namely, the actomyosin contractility at the cell cortex not only tunes mechanical rigidity, but also renders cells rapidly deformable as manifested in cell division and amoeboid migration (Kelkar, Bohec, and Charras 2020; Paluch, Aspalter, and Sixt 2016). Thus, it is of critical importance to disentangle the mode of action of actomyosin in order to understand how cells generate force and shape their morphology.

The actomyosin contractility in nonmuscle cells is mainly attributed to the force generated by nonmuscle myosin II (NMII) (Vicente-Manzanares et al. 2009). NMII contains two heavy chains, two essential light chains, and two regulatory light chains (Vicente-Manzanares et al. 2009; Heissler and Manstein 2013). The myosin regulatory light chains (MLCs) are phosphorylated by myosin light chain kinase (MLCK) and Rho-kinase (ROCK), thereby inducing conformational change in NMII and increasing its motor activity (Vicente-Manzanares et al. 2009). Small chemical compounds have been widely used to perturb the actomyosin contractility, such as blebbistatin (an inhibitor for NMII ATPase activity), Y-27632 (a ROCK inhibitor), and ML-7 (an MLCK inhibitor) (Straight et al. 2003; Uehata et al. 1997; Saitoh et al. 1987). While these compounds have allowed researchers to better understand the function of NMII, it is still technically challenging to control their actions at the subcellular resolution because of their rapid diffusion.

To overcome this limitation, recent efforts have been devoted to the development and application of optogenetic tools to manipulate cell signaling related to actomyosin contractility (Krueger et al. 2019). The most popular approach is to control the activity of RhoA, a member of Rho family small GTPases. Light-induced recruitment of RhoGEF triggers RhoA activation, which in turn activates ROCK and inactivates myosin light chain phosphatase (MLCP) (Wagner and Glotzer 2016; Kimura et al. 1996; Valon et al. 2017) (Fig. 1a). These reactions eventually induce myosin light chain phosphorylation, followed by an increase in the actomyosin contractility. It has been reported that local accumulation of RhoGEF by light increases contractile force at the subcellular scale (Wagner and Glotzer 2016; Oakes et al. 2017; Valon et al. 2017). These technologies allow activation of NMII at the equator and the induction of partial constriction in rounded cells in metaphase (Wagner and Glotzer 2016). Valon *et al*. further demonstrated that trapping of overexpressed RhoGEF to the outer membrane of mitochondria resulted in a decrease in actomyosin contractility (Valon et al. 2017). In addition, depletion of PI(4,5)P2 at the plasma membrane by optogenetic membrane translocation of 5-phosphatase OCRL has been shown to modulate cell contractility and inhibit apical constriction during *Drosophila* embryogenesis (Guglielmi et al. 2015). Although many of these tools enhance actomyosin contractility through RhoA or phospholipids, tools that reduce actomyosin contractility below the basal level have not yet been developed.

**Figure 1.**
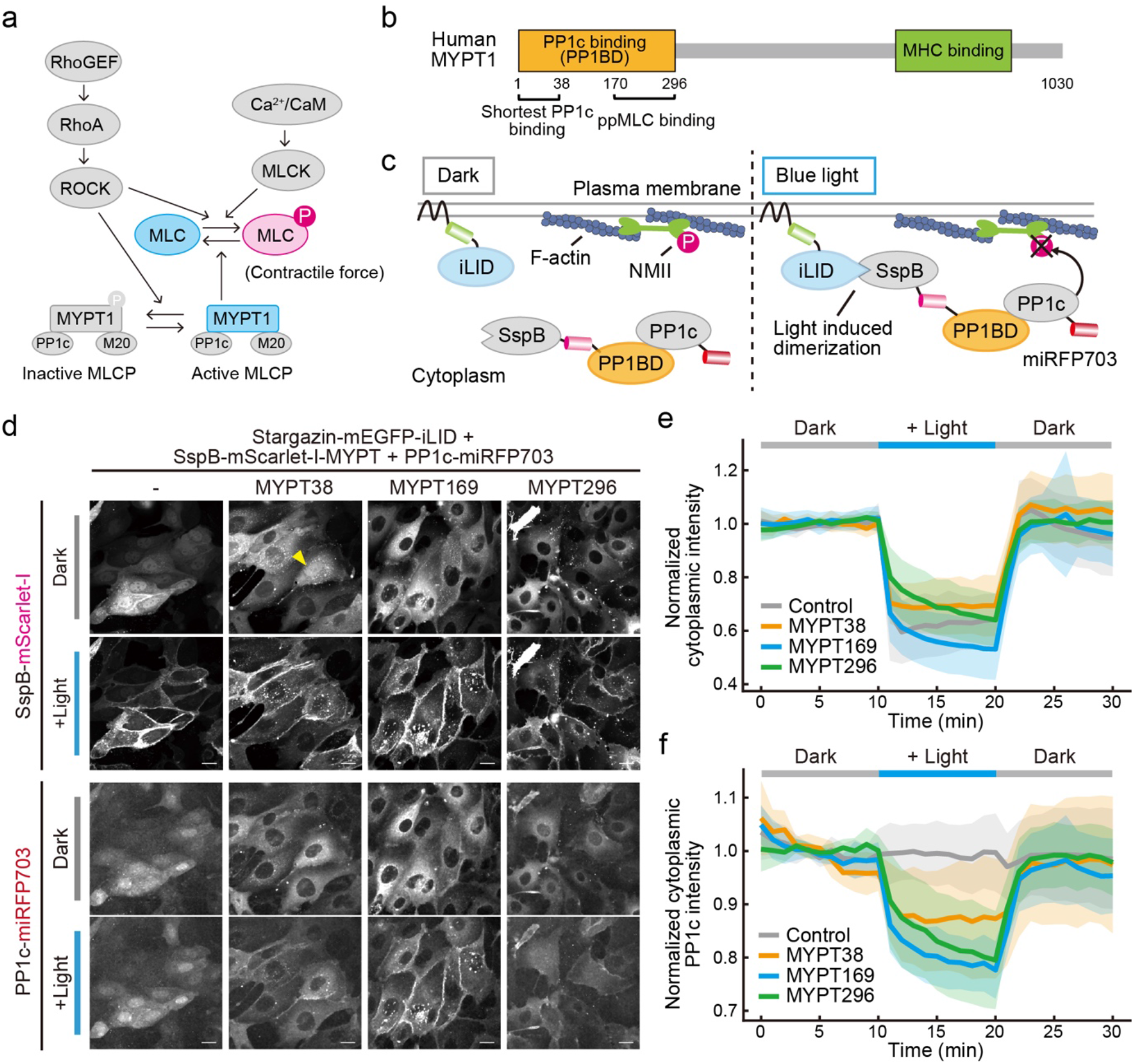
Development of the OptoMYPT system. (a) Schematic illustration of the signaling pathway for the regulation of MLC phosphorylation. (b) Domain structure of human MYPT1. (c) Schematic illustration of the OptoMYPT system. Stargazin-mEGFP-iLID is anchored to the plasma membrane. Upon blue light illumination, the SspB-mScarlet-I-fused PP1c-binding domain (PP1BD) of MYPT1 translocates to the plasma membrane along with endogenous PP1c, and inactivates NMII at the cell cortex. (d) Representative images of the SspB-mScarlet-I or the indicated SspB-mScarlet-I-PP1BDs of MYPT1 (upper two rows) and simultaneously expressing PP1c-miRFP703 (lower two rows) in MDCK cells under the dark condition (the first and third rows) and blue light condition (the second and fourth rows). Stargazin-mEGFP-iLID was also expressed as a localizer in all experiments. The yellow arrowhead indicates a cell showing nuclear accumulation of SspB-mScarlet-I-MYPT38. Scale bar, 20 μm. (e) Quantification of the cytoplasmic fluorescence change in mScarlet-I in panel d. The average values (bold lines) are plotted as a function of time with the SD. n = 15 cells. (f) Quantification of the cytoplasmic fluorescence change in PP1c-miRFP703 in the indicated MDCK cells. The average values (bold lines) are plotted as a function of time with the SD. n = 15 cells.

Here, we report a new optogenetic tool to directly inactivate NMII; the system, called OptoMYPT, is designed to recruit an endogenous catalytic subunit of type Ic phosphatase (PP1c) to the plasma membrane with light, thereby dephosphorylating and inactivating NMII. We demonstrate that MLCs are dephosphorylated and the traction force exerted by cells is reduced at the local area where blue light was illuminated. Moreover, this system was applied to the mechanics of cytokinesis to understand how and to what extent actomyosin-based cortical tension antagonizes contractile ring tension and contributes to the cleavage furrow ingression rate.

## RESULTS

### Design of an OptoMYPT system for reducing intracellular contractile force

To manipulate the intracellular contractile force, we focused on MLCP, which is composed of three subunits, a catalytic subunit PP1c, a regulatory subunit (MYPT1), and a smaller subunit of 20-kDa (M20) (Ito et al. 2004). MYPT1 contains a PP1c-binding domain (PP1BD) and myosin heavy chain (MHC)-binding domain (Fig. 1b). MYPT1 holds PP1c through the PP1BD, and recruits it to NMII to dephosphorylate MLC, leading to the inactivation of NMII. Phosphorylated MLC is mainly localized near the plasma membrane such as at cortical actin and stress fibers, where the NMII exerts mechanical force (Du and Frohman 2009).

Our strategy for the reduction of contractile force is based on inducing membrane translocation of the PP1BD in MYPT1 with light, resulting in the co-recruitment of endogenous PP1c at the plasma membrane and dephosphorylation of MLC. We refer to this system as the OptoMYPT system. It has been reported that the 1 to 38 amino acids (a.a.) in the PP1BD are particularly important for binding to PP1c, and that the 170 to 296 a.a. in the PP1BD serve as a phosphorylated MLC-binding domain (Hirano, Phan, and Hartshorne 1997). As an optogenetic switch in this study, we mainly employed the improved Light-Induced Dimer (iLID) system, which binds to its binding partner, SspB, upon blue light illumination and dissociates from SspB under the dark condition (Guntas et al. 2015). The iLID-based OptoMYPT system consists of a light-switchable plasma membrane localizer, Stargazin-mEGFP-iLID, and an actuator, SspB-mScarlet-I- PP1BD, which is translocated to the plasma membrane for the co-recruitment of the endogenous PP1c with blue light (Fig. 1c). The Stargazin-mEGFP-iLID is suited for the subcellular protein recruitment, because the large N-terminal transmembrane anchor limits the diffusion of SspB proteins (Natwick and Collins 2021). Alternatively, we developed a cryptochrome 2 (CRY2)-based OptoMYPT system, in which CRY2-mCherry-PP1BD was recruited to the plasma membrane with blue light through binding to the plasma membrane localizer CIBN-EGFP-KRasCT (Kennedy et al. 2010).

We first compared the efficacy of light-induced membrane translocation between three different lengths of PP1BDs: 1-38, 1-169, and 1-296 (hereafter referred to as MYPT38, MYPT169, and MYPT296, respectively). In line with the previous study (Wu et al. 2005), SspB-mScarlet-I-PP1BDs accumulated at the nucleus (Fig. S1a,b). To circumvent this problem, the nuclear export signal (NES) was fused with the C terminus of the PP1BDs to export them to the cytoplasm (Fig. S1a,b). As a control, we confirmed that Madin-Darby Canine Kidney (MDCK) cells exhibited the translocation of SspB-mScarlet-I from the cytoplasm to the plasma membrane upon blue light illumination (Fig. 1d,e). SspB-mScarlet-I-MYPT169 showed the best membrane translocation in three differential lengths of PP1BDs (Fig. 1e; Movie S1). We recognized that a small fraction of SspB-mScarlet-I-MYPT38 still resided in the nucleus (Fig. 1d, yellow arrowhead), and CRY2-mCherry-MYPT38 formed aggregates and puncta in a blue light-dependent manner for an unknown reason (Fig. S2). Next, we investigated whether PP1BDs of MYPT1 indeed bind to PP1c and recruit it to the plasma membrane. In control cells, PP1c fused with miRFP703 (PP1c-miRFP703), which was mainly localized at the nucleus, did not show any change in the subcellular localization upon blue light illumination (Fig. 1d,f). As expected, SspB-mScarlet-I-MYPT38, -MYPT169, and -MYPT296 demonstrated similar levels of translocation of PP1c-miRFP703 from the cytoplasm to the plasma membrane upon blue light illumination (Fig. 1d,f; Movie S2). We further evaluated the effect of overexpression of PP1BD on basal phosphorylation of MLC (ppMLC) by western blotting analysis. Interestingly, MLCs were almost completely dephosphorylated in MYPT296-overexpressing cells (Fig. S1c). Indeed, the cells expressing MYPT296 showed flattened morphology with membrane protrusions similar to the morphology of cells treated with ROCK inhibitor or myosin inhibitor (Worthylake and Burridge 2003; Totsukawa et al. 2004) (Fig. S1a). This result is consistent with the previous report pointing out the existence of the ppMLC-binding domain at the 170–296 a.a., which may facilitate recruitment of PP1BD to NMII and dephosphorylation of MLC without any stimulation (Hirano, Phan, and Hartshorne 1997). Taken together, these results led us to conclude that MYPT169 is well suited for the OptoMYPT system.

### Characterization of the OptoMYPT system

To evaluate whether the OptoMYPT dephosphorylates ppMLC in a blue light-dependent manner, we directly stained ppMLC with immunofluorescence. The blue light was locally illuminated for 15 min, followed by fixation and immunofluorescence staining with the anti-ppMLC antibody (Fig. 2a). We herein adopted a CRY2-based OptoMYPT system (Kennedy et al. 2010), because the slower dissociation kinetics of the CRY2-CIB system compared to that of the iLID-SspB system was preferable for this experiment. Local illumination of blue light induced spatially restricted recruitment of CRY2-mCherry and CRY2-mCherry-MYPT169 (Fig. 2b, right column). In addition, the local recruitment of CRY2-mCherry-MYPT169, but not CRY2-mCherry, attenuated the ppMLC signal (Fig. 2b, left column). The quantification of ppMLC fluorescence intensity in dark and light illuminated areas (Fig. 2a) revealed a significant reduction in the ppMLC level (Fig. 2c).

**Figure 2.**
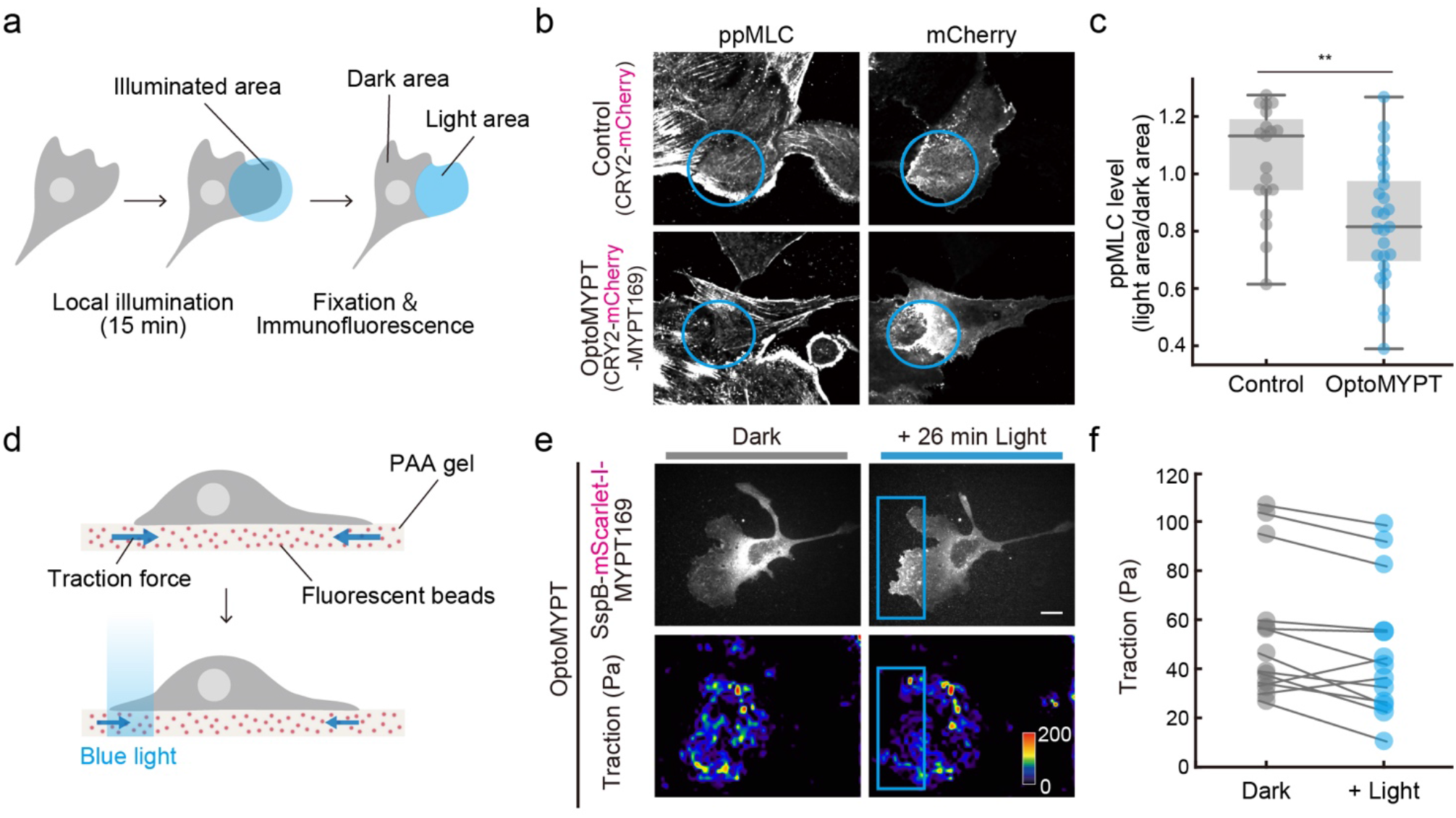
Characterization of the OptoMYPT system. (a) Schematic illustration of an experimental procedure to quantify ppMLC levels. The blue light was locally focused on the lamellipodial area in each cell, followed by fixation and immunostaining. (b) Immunofluorescence analysis of ppMLC after local blue-light illumination. The upper and lower images show MDCK cells expressing CRY2-mCherry and CRY2-mCherry-MYPT169, respectively. CIBN-EGFP-KRasCT was also expressed as a localizer in both experiments. Blue circles indicate the area illuminated with blue light. Scale bar, 10 μm. (c) The ppMLC level was quantified by dividing the mean fluorescence intensity of the light area by that of the dark area in panel a, and shown as a box plot, in which the box extends from the first to the third quartile with the whiskers denoting 1.5 times the interquartile range. n = 19 and 27 cells for the control and OptoMYPT, respectively. \*\**p* < 0.01 (student’s *t*-test). (d) Schematic illustration of the traction force microscopy. (e) Traction force measurement in an MDCK cell expressing SspB-mScarlet-I-MYPT169 and Stargazin-mEGFP-iLID. Blue rectangles indicate blue-light illuminated areas. Traction force (Pa) is represented as a pseudo color. Scale bar, 20 μm. (f) Quantification of the traction force before and after blue-light illumination. n = 13 cells.

The decrease in ppMLC by OptoMYPT prompted us to examine the effect on contractile force by light. To do this, traction force microscopy was applied to randomly migrating MDCK cells expressing the OptoMYPT system. The cells were seeded on polyacrylamide gel containing infra-red fluorescence beads, so that we could infer the traction force by the displacement of fluorescence beads and the mechanical properties of the polyacrylamide gel (Fig. 2d). The blue light was locally focused on the lamellipodial region, where the cells were generating strong traction force. SspB-mScarlet-I-MYPT169 was successfully recruited to the locally illuminated area (Fig. 2e, upper). Under this condition, the traction force was reduced after blue light illumination (Fig. 2e, lower; Fig. 2f, Movie S3). These results indicate that the OptoMYPT system can dephosphorylate ppMLCs by local blue light illumination, leading to a reduction of traction force.

### Membrane protrusion induced by local dephosphorylation of ppMLC with OptoMYPT

We next examined whether local dephosphorylation of ppMLC and reduction of contractile force have an impact on cell morphology by using OptoMYPT. The control MDCK cells that expressed SspB-mScarlet-I, Stargazin-mEGFP-iLID, and Lifeact-miRFP703 demonstrated local accumulation of SspB-mScarlet-I by blue light illumination, but did not show morphological change (Fig. 3a). On the other hand, the OptoMYPT-expressing MDCK cells reproducibly showed peripheral membrane protrusions in the blue-light exposed area (Fig. 3b; Movie S4). We evaluated light-induced membrane protrusion with a kymograph and time-course graph, which showed the movement of the cell edge upon blue light illumination in OptoMYPT-expressing cells (Fig. 3c, d). The protruding membrane was subsequently maintained under dark conditions. Interestingly, we often recognized membrane retraction on the opposite side of the blue-light illumination area (Fig. 3e, arrowhead; Movie S5). The OptoMYPT-induced membrane protrusion and retraction are consistent with the previous reports showing Rac1 activation by inhibiting NMII or ROCK (Even-Ram et al. 2007; Martin et al. 2016) and regulation of cell polarity by membrane tension (Houk et al. 2012).

**Figure 3.**
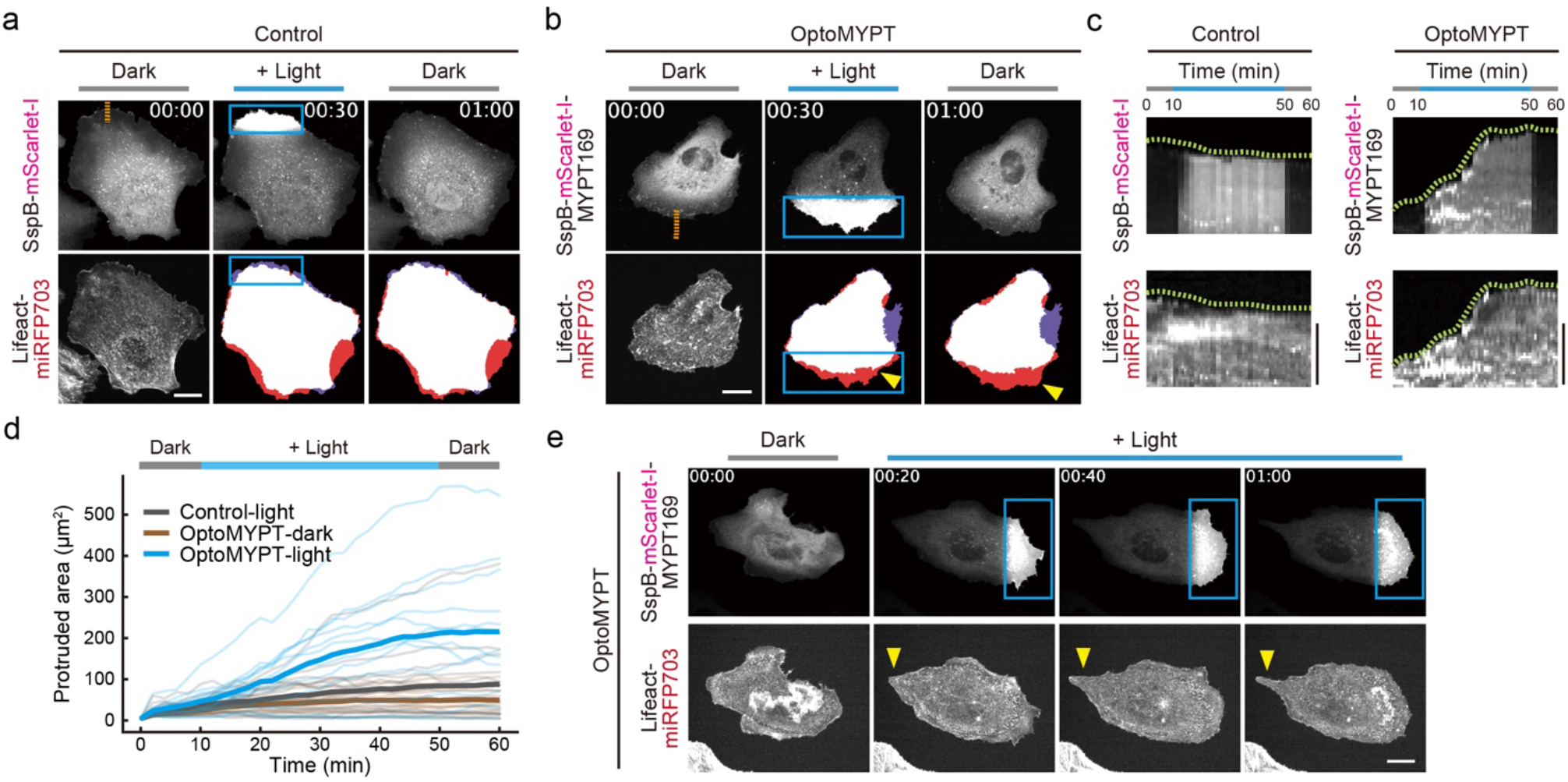
Induction of membrane protrusion by the OptoMYPT system. (a, b) Simultaneous visualization of SspB-mScarlet-I (a) or SspB-mScarlet-I-MYPT169 (b) with F-actin (Lifeact-miRFP703) in MDCK cells. Blue rectangles indicate blue-light illuminated areas. The middle and right bottom images show binary images reconstructed from Lifeact-miRFP703 images; red and purple areas represent protruding and retracting areas, respectively. The yellow arrowheads depict membrane protrusion. Scale bar, 20 μm. (c) Kymographs were drawn along the orange dashed lines in panels a and b. Green dashed lines show cell boundaries. Scale bar, 5 μm. (d) Quantification of the local protruding areas under the indicated conditions. The total protruded area was calculated by subtracting the cell area in the locally illuminated region at t = 0 from that at each time point. Local blue light was illuminated from 10 to 50 min. The thin and bold lines indicate the individual and averaged data, respectively. n = 15, 13, and 12 cells for Control-light (local blue light illumination), OptoMYPT-dark (dark condition), OptoMYPT-light (local blue light illumination), respectively. (e) Representative images of the induction of membrane retraction (yellow arrowhead) on the opposite side of the blue-light illuminated area.

### Acceleration of the ingression rate of cleavage furrows during cytokinesis by the optical relaxation of cortical tension

We next applied the OptoMYPT system to elucidate the mechanical regulation of the actin cortex during cytokinesis. In this process, a contractile ring, which is mainly composed of F-actin and NMII, transiently forms in the equatorial plane and generates tension to constrict the ring and divide a cell into two daughter cells (Fig. 4a, solid arrows). The tension in the contractile ring is counteracted by the tension generated by cortical actomyosin (Fig. 4a, dashed arrows). Recent studies showed that genetic and pharmacological perturbations to the cortical actomyosin disrupted proper cytokinesis (Taneja et al. 2020; Yamamoto et al. 2019; Sedzinski et al. 2011; O’Connell, Warner, and Wang 2001), suggesting that the cortical tension plays a vital role. However, there is still controversy in regard to the importance of cortical tension, because it is difficult to estimate the strength of cortical tension relative to the contractile ring.

**Figure 4.**
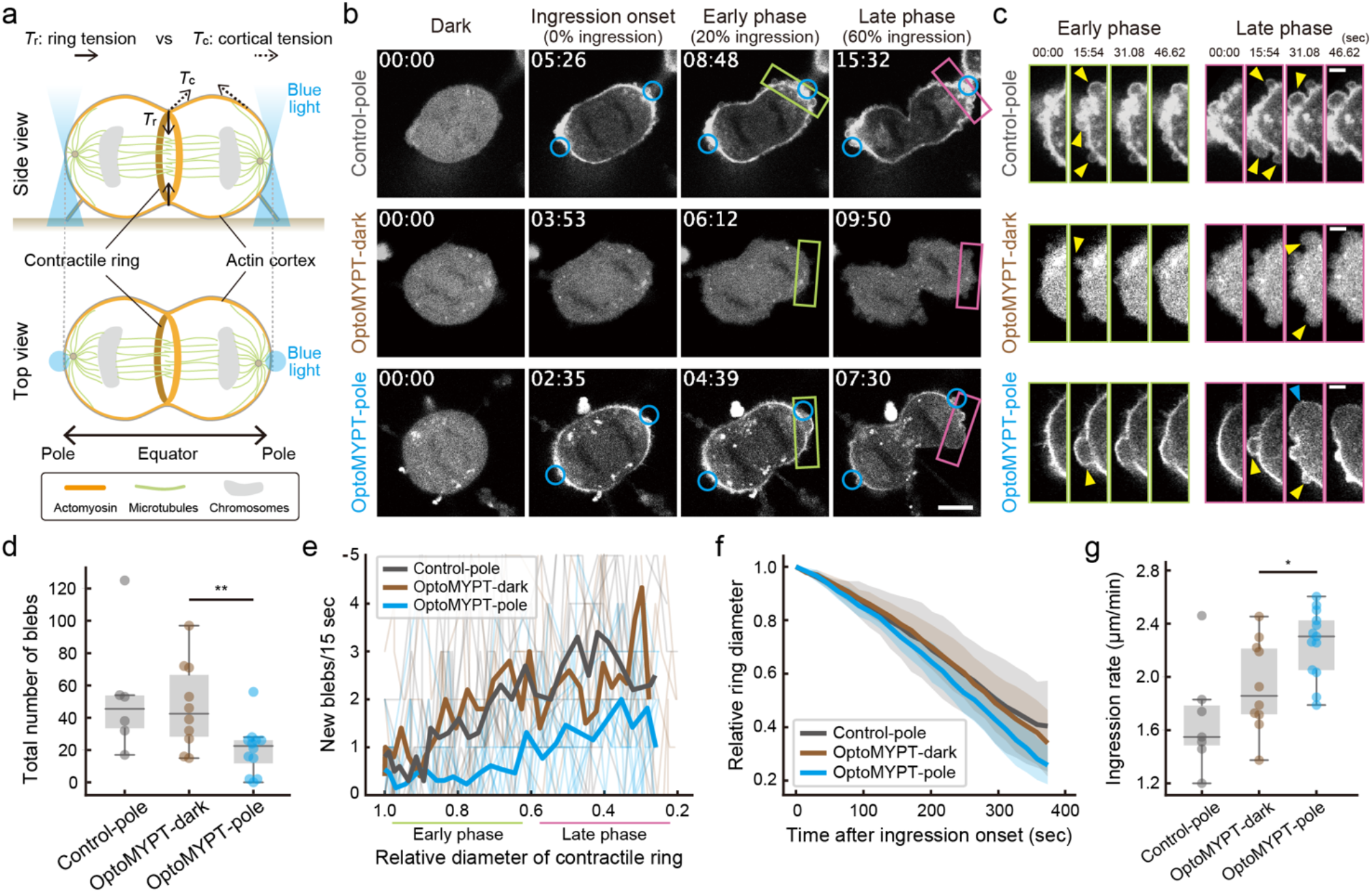
Examination of the actomyosin-based cortical tension during cytokinesis with OptoMYPT. (a) Schematic illustration of cytokinesis in animal cells. Solid and dashed arrows indicate ring tension and cortical tension, respectively. Orange, green, and gray objects indicate actomyosin, microtubules, and chromosomes, respectively. Blue light is focused on the poles on both sides. (b) Representative images of SspB-mScarlet-I (upper) or SspB-mScarlet-I-MYPT169 (middle and lower) in MDCK cells during cytokinesis. Blue circles in the upper and lower panels indicate blue-light illuminated areas. Middle panels represent cytokinesis of a cell under dark condition. Scale bar, 10 μm. (c) Inset images of polar blebbing (the green and magenta boxed regions in panel b, representing the early and late phases, respectively). Yellow and blue arrowheads indicate small and large new blebs per stack, respectively. Scale bar, 3 μm. (d, e) Quantification of the total number of blebs during cytokinesis, shown as a box plot (d), and of the number of new blebs emerged within 15.54 sec, shown as a line graph (e), in which thin and bold lines indicate individual and averaged data, respectively. n = 5, 10, and 13 cells for Control-pole, OptoMYPT-dark, and OptoMYPT-pole, respectively. \*\**p* < 0.01 (student’s *t*-test). (f, g) Quantification of the furrow ingression rate after ingression onset. Averaged relative diameters are plotted as a function of time with the SD (f). The ingression rate was estimated by calculating the slope of the ingression rate from 1.0 to 0.6 in panel f, and shown as a box plot (g). n = 7, 10, and 13 cells for Control-pole, OptoMYPT-dark, and OptoMYPT-pole, respectively. \**p* < 0.05 (student’s *t*-test).

To address this issue, we perturbed the cortical tension by using the OptoMYPT system. Blue light was locally and repeatedly illuminated at both poles from the onset of chromosome segregation (Fig. 4a). SspB-mScarlet-I and SspB-mScarlet-I-MYPT169 were trapped in the polar region (Control-pole and OptoMYPT-pole, respectively) (Fig. 4b; Movies S6, S7). Because the overexpression of OptoMYPT might reduce basal actomyosin activity (Fig. S1c), cells expressing SspB-mScarlet-I-MYPT169 under a dark condition throughout cytokinesis (OptoMYPT-dark) were also used as a control (Fig. 4b; Movie S8).

To evaluate the reduction of cortical tension by local activation of the OptoMYPT, we focused on the dynamics of membrane blebbing during cytokinesis (Fig. 4c). Because it has been reported that high cortical tension causes membrane blebbing (Tinevez et al. 2009), we presumed that the appearance of membrane blebbing and the number of blebs could be used as an indicator of cortical tension. We counted the number of membrane blebs since the onset of cleavage furrow ingression, and found a decrease in the number of blebs in OptoMYPT-pole cells as compared to Control-pole and OptoMYPT-dark cells (Fig. 4c, d). This result is consistent with previous observations, in which NMIIA-knockout or knockdown alleviated cortical tension during cytokinesis (Yamamoto et al. 2019; Taneja et al. 2020). Furthermore, smaller membrane blebbing emerged in Control-pole and OptoMYPT-dark cells from the early phase of cleavage furrow ingression, and the number of blebs gradually increased as furrow ingressed, whereas OptoMYPT-pole cells exhibited larger membrane blebbing from the late phase of cleavage furrow ingression (Figs. 4e and S3). Based on these results, we concluded that the cortical tension in both poles decreased through local application of the OptoMYPT system.

Finally, to estimate the strength of cortical tension relative to that of contractile ring tension, we adapted a coarse-grained physical model describing the mechanics of cytokinesis (Sedzinski et al. 2011; Yoneda and Dan 1972; Turlier et al. 2014). In this model, the furrow ingression rate (*v*) is considered to be proportional to the difference between the contractile force of the ring (*F*_r_) and the resisting force exerted by the cortices (*F*_c_), *ν* ∝ *F*_r_ - *F*_c_ (see the Materials and Methods, Fig. S4, and the Supplementary Discussion). Thus, we measured the furrow ingression rate under each condition. The ingression rate of the cleavage furrow was significantly higher in OptoMYPT-pole cells (2.26 ± 0.25 μm/min) than in Control-pole cells (1.55 ± 0.20 μm/min) and OptoMYPT-dark cells (1.93 ± 0.33 μm/min) (Fig. 4f, g), indicating that the reduced cortical tension accelerates the cleavage furrow ingression rate. These results highlight the negative contribution of the cortical tension to the cleavage furrow ingression. Based on the coarse-grained physical model and experimental data, we estimated that the cortical tension corresponds to at least 14.6% of the ring tension during cytokinesis (see the Supplementary Discussion).

## DISCUSSION

In this study, we developed a new optogenetic tool, OptoMYPT, and demonstrated light-dependent relaxation of cellular forces at the subcellular level. The OptoMYPT is substantially different from existing optogenetic tools related to the cell mechanics in two ways. First, the OptoMYPT reduces contractile forces below the basal level, and therefore provides additional flexibility for *in situ* control of actomyosin contractility and cellular morphology. Second, the OptoMYPT directly regulates NMII through MLC dephosphorylation, whereas optogenetic modulation of RhoA activity or PI(4,5)P2 may affect pathways other than NMII, since RhoA and PI(4,5)P2 are known to control various downstream effectors such as ROCK, mDia, and the ezrin-radixin-moesin (ERM) proteins (Oshiro, Fukata, and Kaibuchi 1998; Yonemura et al. 2002; Narumiya, Tanji, and Ishizaki 2009).

Using the OptoMYPT system, we experimentally revealed the negative contribution of the cortical tension to the cleavage furrow ingression rate during cytokinesis; *i.e*., the decrease in cortical tension at both poles by OptoMYPT accelerates furrow ingression (Figs. 4 and 5). It has been reported that the reduction of cortical tension by laser ablation in the polar region decelerates cleavage furrow ingression in cytokinesis of *C. elegans* embryos (Khaliullin et al. 2018). This discrepancy could be due to the difference in the force balance between the pole and equator; in *C. elegans* embryos, NMII is actively removed from the polar region and accumulates at the equator due to the cortical flow, and thus cortical tension is much weaker than ring tension (Chapa-Y-Lazo et al. 2020; Reymann et al. 2016). Meanwhile, our results indicate that cortical tension in cultured mammalian cells is comparable to contractile ring tension (Fig. 4), which is in good agreement with the previous work (Sedzinski et al. 2011). Such a high cortical tension is advantageous because it confers shape stability to mitotic cells (Sedzinski et al. 2011). This has been corroborated in a recent paper demonstrating that high cortical stiffness in cancer cells allows them to divide in a confined environment (Matthews et al. 2020). The benefit of cortical tension is not required in *C. elegans* embryos, because they are covered and protected by a rigid eggshell. However, high cortical tension is a double-edged sword, because excessive cortical tension induces cytokinetic shape oscillation and abscission failure (Mukhina, Wang, and Murata-Hori 2007; Taneja et al. 2020; Girard et al. 2004). The estimated cortical tension relative to ring tension (*F*_c_/*F*_r_), ~14.6%, may achieve a balance between morphological maintenance and timely cytokinesis.

**Figure 5.**
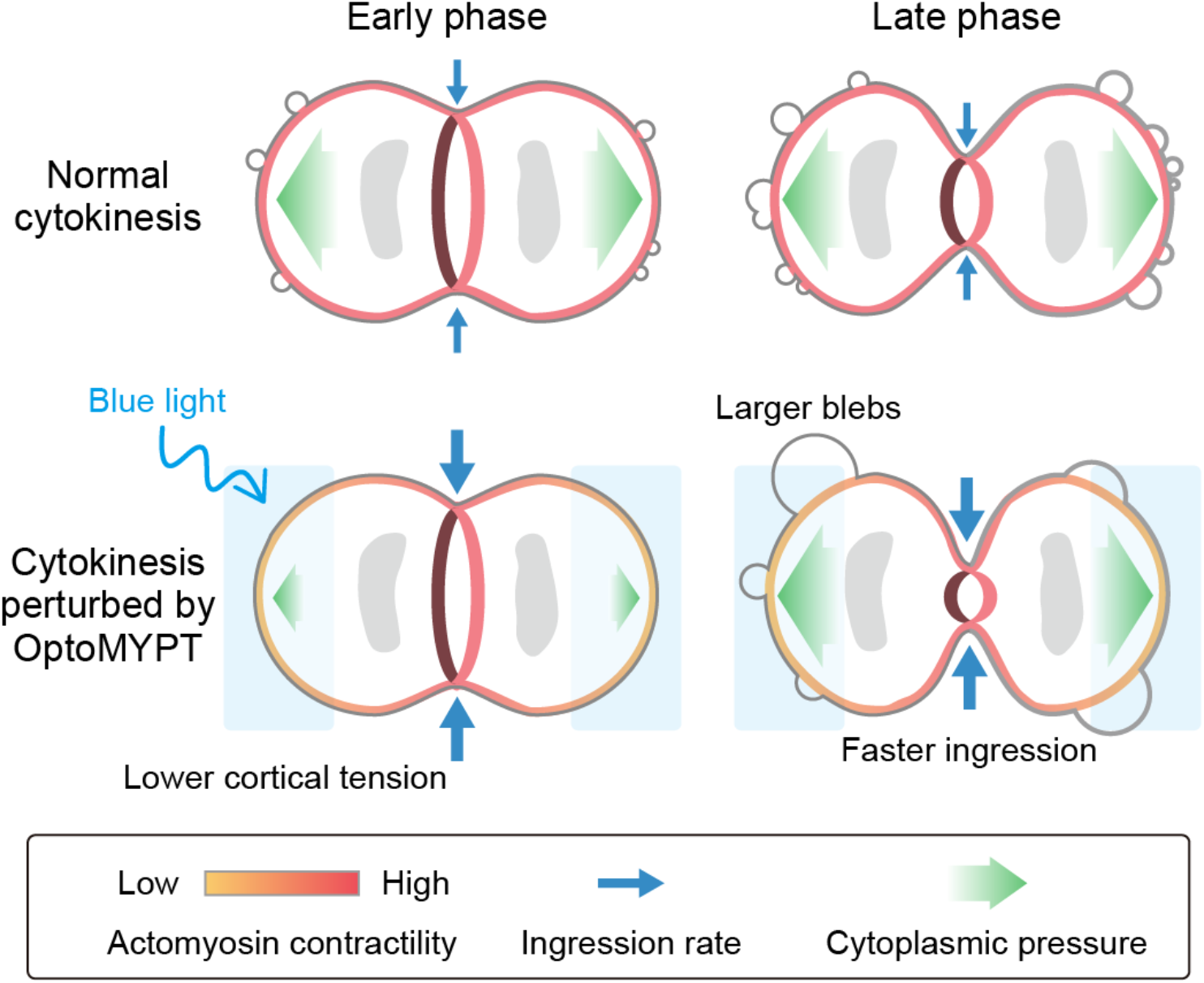
Schematic model of mechanical regulation of the contractile ring and the actin cortex during cytokinesis. In control cells, high cortical tension acts as a decelerator of the ring constriction. The high cortical tension and cytoplasmic pressure induce blebbing from the early phase of cytokinesis. The increased cytoplasmic pressure associated with ring constriction in the late phase is released by the increased number of blebs. In pole-illuminated OptoMYPT cells, the cleavage furrow ingression is accelerated due to the decrease in the cortical tension. As cleavage furrow ingression progresses, large blebs emerge due to the increased cytoplasmic pressure.

We also focused on the dynamics of membrane blebbing during cytokinesis (Fig. 4). Although the functional significance of blebbing is poorly understood, except in relation to cell migration, the mechanisms underlying membrane blebbing have been increasingly reported. Membrane blebbing is initiated by local rupture of the actin cortex and/or detachment of the actin cortex from the plasma membrane (Charras and Paluch 2008). The growth of membrane blebs requires a condition under which hydraulic pressure caused by actomyosin-based cortical tension overcomes plasma membrane tension (Tinevez et al. 2009). We showed that numerous small blebs emerged in the “early phase” of furrow ingression in control MDCK cells, and these were suppressed by relaxation of cortical tension with OptoMYPT (Figs. 4e, 5, and S3). This phenomenon is clearly consistent with the aforementioned mechanisms. Meanwhile, MDCK cells expressing OptoMYPT exhibited large blebs in the “late phase” of cleavage furrow ingression when both poles were illuminated with blue light (Figs. 4 and 5). The late onset of blebbing in the pole-illuminated cells probably occurs because the progression of cleavage furrow ingression raises hydraulic pressure, and thereby induces detachment of the actin cortex from the plasma membrane, leading to the pressure release through bleb formation. To the best of our knowledge, the mechanics of bleb formation in the “late phase” of the furrow ingression are a hitherto unrecognized process.

There still remain some issues to be addressed with respect to the OptoMYPT. First is the issue of substrate specificity in OptoMYPT. We could not exclude the possibility that the OptoMYPT dephosphorylates additional substrates other than MLC. However, based on the fact that the OptoMYPT activation at both poles during cytokinesis did not induce bleb formation, it is unlikely that OptoMYPT dephosphorylates and inactivates the ERM proteins cross-linking the plasma membrane and actin cortex to prevent membrane bleb formation (Rodrigues et al. 2015). The next issue is the localizer of optogenetic switches. In the current OptoMYPT system, dephosphorylation of MLC was induced by recruiting MYPT169 to the plasma membrane upon illumination with blue light. However, it is plausible that the OptoMYPT dephosphorylates and inactivates only the NMII existing in the vicinity of the plasma membrane, and not the NMII in the other subcellular locations. The use of a localizer that is closer to endogenous active NMII or a specific localization such as the apical membrane of epithelial cells could result in an OptoMYPT system with better specificity and spatial resolution than the current version.

The OptoMYPT system will provide opportunities not only to understand the mechanics of morphogenesis, but also to shape the morphology of cells and tissues with precision and flexibility as desired. Recent papers have applied optogenetic systems *in vivo*, and succeeded in inducing arbitrary forms of the apical constriction (Izquierdo, Quinkler, and De Renzis 2018). By combining red light-responsive optogenetic tools such as PhyB-PIF with blue light-responsive tools (Levskaya et al. 2009; Uda et al. 2017), it will be possible to create more sophisticated morphology with an increase or decrease in contractile force in the same cells and tissues.

## Materials and Methods

### Plasmids

The cDNAs of human MYPT1 and PP1c were derived from HeLa cells (Human Science Research Resources Bank). SspB-mScarlet-I-PP1BDs were obtained by Gibson assembly cloning, combining the SspB obtained from pPBbsr tgRFPt-SspB and cDNAs of PP1BDs. Stargazin-mEGFP-iLID was obtained by Gibson assembly cloning, combining the Stargazin derived from Stargazin-GFP-LOVpep (plasmid #80406: Addgene) and pPBbsr Venus-iLID-CAAX. CRY2 and CIBN-EGFP-KRasCT were obtained from pCX4puro-CRY2-CRaf and pCX4neo-CIBN-EGFP-KRasCT (Aoki et al. 2013) and inserted into the pCAGGS vector (Niwa, Yamamura, and Miyazaki 1991). The cDNAs of Lifeact and NES were obtained by oligo DNA annealing and ligation, and inserted into each vector. The cDNA of hyPBase, an improved PiggyBac transposase (Yusa et al. 2011), was synthesized (FASMAC), and inserted into the pCAGGS vector. The nucleotide sequences of newly generated constructs are provided in Table S1.

### Cell culture

MDCK cells (no. RCB0995: RIKEN Bioresource Center) were maintained in minimal essential medium (MEM; 10370-021: ThermoFisher Scientific) supplemented with 10% fetal bovine serum (FBS; 172012-500ML: Sigma), 1x Glutamax (35050-061: ThermoFisher), and 1 mM sodium pyruvate (11360070: ThermoFisher) in a 5% CO_2_ humidified incubator at 37°C. The cells were split as previously reported (Aoki et al. 2017).

### Transfection

Because PP1BDs of MYPT1 fused with fluorescent proteins tend to aggregate for long-term expression, most experiments were performed by transient expression. The MDCK cells were electroporated by using Nucleofector IIb (Lonza) according to the manufacturers’ instructions (T-023 program) with a house-made DNA- and cell-suspension solution (4 mM KCl, 10 mM MgCl_2_, 107 mM Na_2_HPO_4_, 13 mM NaH_2_PO_4_, 11 mM HEPES pH. 7.75). After electroporation, the cells were plated on collagen-coated 35-mm glass-base dishes.

### Establishment of stable cell lines

For transposon-mediated gene transfer, MDCK cells were transfected with PiggyBac donor vectors and PiggyBac transposase-expressing vectors at a ratio of 3:1. One day after transfections, cells were treated with 10 μg/mL blasticidin S (InvivoGen, San Diego, CA) or 1.0 μg/mL puromycin (InvivoGen) for selection. The bright bulk cell population was collected using a cell sorter (MA900; SONY).

### Live-cell fluorescence imaging

Cells were imaged with an IX83 inverted microscope (Olympus, Tokyo) equipped with an sCMOS camera (Prime: Photometrics, Tucson, AZ; or ORCA-Fusion BT: Hamamatsu Photonics, Hamamatsu, Japan), a spinning disk confocal unit (CSU-W1; Yokogawa Electric Corporation, Tokyo), and diode lasers at wavelengths of 488 nm, 561 nm, and 640 nm. An oil immersion objective lens (UPLXAPO60XO, N.A. 1.42; Olympus) or an air/dry objective lens (UPLXAPO40X, N.A. 0.95; Olympus) was used. The excitation laser and fluorescence filter settings were as follows: Excitation laser, 488 nm (mEGFP), 561 nm (mScarlet-I), and 640 nm (miRFP703); dichroic mirror, DM 405/488/561 dichroic mirror (mEGFP, mScarlet-I, and miRFP703); emission filters, 500–550 nm (mEGFP), 580–654 nm (mScarlet-I), and 665–705 nm (miRFP703). During observation, cells were incubated with a stage incubator set to 37 °C and containing 5% CO_2_ (STXG-IX3WX; Tokai Hit).

For global illumination of the blue light, blue LEDs (450 nm) were manually illuminated from the top of the stage or pulsed blue light (488 nm) was illuminated through the objective lens. For local light illumination in the interphase cells, a digital micromirror device (Polygon 400; Mightex) mounted on the IX83 microscopic system, and pT-100 (CoolLED) were used. For local light illumination during cytokinesis, an SP8 FALCON inverted confocal laser scanning microscope (Leica) equipped with a water immersion objective lens (HC PL APO 63x/1.20 W motCORR; Leica) was used. Local light illumination was started using the FRAP function just after chromosome segregation onset. We illuminated every 3.11 sec, and acquired images every 15.54 sec. The positions of regions of interest (ROIs) were manually corrected every 2 min in all samples.

For all time-lapse imaging, MDCK cells were plated on 35 mm glass-base dishes (IWAKI). Before time-lapse imaging, the medium was replaced with FluoroBrite (Invitrogen) supplemented with 10% FBS, 1x Glutamax.

### Immunofluorescence

Cells were fixed with 3.7% formaldehyde in PBS for 20 min, followed by permeabilization by 5 min incubation in 0.05% Triton X-100-containing PBS. Samples were soaked for 30 min in Can Get Signal immunostain (solution A) (Toyobo, Japan), and then incubated with ppMLC antibody (1:50 dilution; Cell Signaling Technology #3674) in Can Get Signal immunostain (solution A) for 1 h at room temperature. Next, the cells were washed 3 times with PBS, and then incubated for 1 h at room temperature with Alexa 555-conjugated anti-rabbit IgG (1:100 dilution; ThermoFisher) in Can Get Signal immunostain (solution A). Finally, the cells were washed 3 times with PBS and subjected to fluorescence imaging.

### Traction force microscopy

Polyacrylamide gel substrates were prepared in accordance with previously published protocols (Tambe et al., 2011; Trepat et al., 2009). In brief, the gel solution was prepared with 4% acrylamide, 0.1% bisacrylamide, 0.8% ammonium persulfate, 0.08% TEMED (Nacalai Tesque), and 5% deep red fluorescent carboxylate-modified beads (0.2 μm diameter; F8810; Thermo Fisher Scientific). 13 μL of the mixture was added to a 35 mm glass-base dish (IWAKI) and then covered with a glass coverslip of 15 mm diameter (Matsunami). After gel polymerization at room temperature, the surface was coated with 0.3 mg/mL type I collagen (Nitta Gelatin, Osaka, Japan) using 4 mM sulphosuccinimidyl-6-(4-azido-2-nitrophenylamino) hexanoate (Sulfo-SANPAH; Pierce). Cells were seeded on the gel, and imaged with a spinning disk confocal microscope. To quantify the traction force, two Fiji plugins, *i.e*., the iterative PIV and FTTC plugins, were used.

Note that Young’s modulus of the gel was estimated as ~2 kPa according to a previous report (Tse and Engler 2010). The traction force in locally illuminated areas was used for the quantification.

### Immunoblotting

Cells were lysed in 1x SDS sample buffer. After sonication, the samples were separated by 5–20% gradient SDS-polyacrylamide gel electrophoresis (Nagaiki precast gels; Oriental Instruments, Ltd.) and transferred to polyvinylidene difluoride membranes (Millipore). After blocking with Odyssey Blocking Buffer-TBS (LICOR Biosciences) for 1 h, the membranes were incubated with primary antibodies overnight at 4°C, followed by the secondary antibodies for 1 h at room temperature. For primary antibodies, ppMLC antibody (1:500 dilution; Cell Signaling Technology #3674), phospho-Ezrin/Radixin/Moesin antibody (1:500 dilution, Cell Signaling Technology #3276), and α-Tubulin antibody (DM1A) (1:5000 dilution; sc-32293: Santa Cruz Biotechnology) were diluted in Odyssey Blocking Buffer-TBS. For secondary antibodies, IRDye680LT-conjugated goat polyclonal anti-rabbit IgG (H + L) (1:5000 dilution; LI-COR Bioscience) and IRDye800CW-conjugated donkey polyclonal anti-mouse IgG (H + L) (1:5000 dilution; LI-COR Bioscience) were diluted in Odyssey Blocking Buffer-TBS. Proteins were detected with an Odyssey infrared scanner (LI-COR Bioscience).

### Imaging analysis

All fluorescence imaging data were analyzed and quantified by Fiji (Image J). For all images, the background was subtracted and images were registered by StackReg, a Fiji plugin to correct misregistration, if needed. To quantify the cytoplasmic fluorescence intensity changes in Figure 1, the ROI was selected in each image and normalized by the mean fluorescence intensity of the first 10 images under the dark condition. To quantify the area of membrane protrusion in Figure 3, the ROI was chosen so as to coincide with the local light-irradiated area. In OptoMYPT-dark cells, the ROI was a region of lamellipodia similar to that of light-illuminated cells. Fluorescence images of Lifeact-miRFP703 were binarized and the difference between the area at each time point and at t = 0 was calculated.

### Physical modeling

According to previous modeling efforts on the force balance in a dividing cell (Sedzinski et al. 2011; Yoneda and Dan 1972), the temporal change in radius of the contractile ring, *R_r_*, is described as

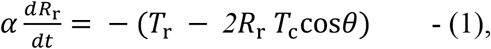

where *α* is ring viscosity, *T*_r_ is the tension of the contractile ring, *T*_c_ is the tension in the actomyosin cortex, and *θ* is the angle between the equatorial plane and polar surface at the furrow (see Fig. S4 for the details). Note that our model was simplified from Eq. S1 in the report of (Sedzinski et al. 2011), since we consider symmetric pole shapes. For brevity, let *F*_r_ and *F*_c_ denote *T*_r_ (ring tension) and *2R*_r_*T*_c_ cos *θ* (cortical tension), respectively (Fig. S4). Then, the furrow ingression rate, *v*, can be expressed as

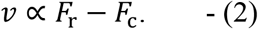

## Acknowledgments

We thank all members of the Aoki Laboratory for their helpful discussions and assistance. We also thank Michiyuki Matsuda, Naoya Hino (Kyoto University), and Hisayo Fukuda (Kansai Medical University) for the advice on electroporation. K.Y. was supported by a JSPS KAKENHI Grant (19J20538). Y.K. was supported by JSPS KAKENHI Grants (19K16207, 19H05675). K.A. was supported by a CREST, JST Grant (JPMJCR1654), and JSPS KAKENHI Grants (18H0244, 19H05798). This research was supported by Joint Research of the Exploratory Research Center on Life and Living Systems (ExCELLS) (ExCELLS program No.18-204, 19-205, 20-204).

## Author contributions

K.Y. and K.A. designed the research. K.Y., H.M., M.I. and S.S. performed experiments. K.Y. and H.M. analyzed data. K.Y., Y.K., and K.A. wrote the manuscript.

