## Supplemental Information for "Optogenetic relaxation of actomyosin contractility uncovers mechanistic roles of cortical tension during cytokinesis"

### Supplementary Discussion

#### Physical modeling

Previous studies have established a simple equation for the equilibrium of forces between the contractile ring and the two polar cortices (Yoneda and Dan 1972; Sedzinski et al. 2011; Turlier et al. 2014). Based on the Young-Laplace law, the force balance is written as

$$\sigma_{\mathrm{r}} = 2R_{\mathrm{r}}T_{\mathrm{c}}\cos\theta, \text{ (S1)}$$

where $\sigma_{\mathrm{r}}$ is the net tension in the ring and $T_{\mathrm{c}}$ is the tension in the polar cortices (see Fig. S4 for the definitions of the geometric factors $R_{\mathrm{r}}$ and $\theta$). Since the turnover of cortical actin (~10 sec) is much faster than the time scale of cytokinesis (~10 min), the material property of polar cortices can be described as active viscous liquid (Turlier et al. 2014), and thus the cortical tension depends on the activity of myosin motors at the cortices. On the other hand, the net tension in the ring is composed of two factors, tension generated by myosin motors $T_{\mathrm{r}}$ and viscous resistance by, e.g., cross-linkers, as

$$\sigma_{\mathrm{r}} = T_{\mathrm{r}} + \alpha\frac{dR_{\mathrm{r}}}{dt}, \text{ (S2)}$$

where the coefficient $\alpha$ represents the viscosity of the actin network constituting the ring (Sedzinski et al. 2011). The cortices can be described as an active viscous membrane having tension $T_{\mathrm{c}}$. Here, if we combine Eqs. S1 and S2, we obtain

$$\alpha\frac{dR_{\mathrm{r}}}{dt} = -(T_{\mathrm{r}} - 2R_{\mathrm{r}}T_{\mathrm{c}}\cos\theta), \text{ (S3)}$$

This is Eq. (1) in the main text.

#### Lower boundary of the cortical tension

In our model, the furrow ingression rate of OptoMYPT-dark cells, $v$, can be expressed as

$$v \propto F_{\mathrm{r}} - F_{\mathrm{c}}, \text{ (S4)}$$

where $F_r$ and $F_c$ are the ring tension and the cortical tension, respectively (Fig. S4). This is Eq. (2) in the main text. The furrow ingression rate of OptoMYPT-pole cells, $v'$, can be expressed as

$$v' \propto F_{\mathrm{r}} - F_{\mathrm{c}}', \text{ (S5)}$$

where $F_c'$ is the cortical tension upon blue light illumination. Note that $F_r$ is considered to be constant under each condition, because blue light was locally illuminated to the polar cortices. Taken together with Eqs. S4 and S5, we obtain

$$v/v' = (F_{\mathrm{r}} - F_{\mathrm{c}})/(F_{\mathrm{r}} - F_{\mathrm{c}}') > (F_{\mathrm{r}} - F_{\mathrm{c}})/(F_{\mathrm{r}}). \text{ (S6)}$$

The rightmost expression represents the case where blue light illumination completely reduces cortical tension to zero. We rearrange the above formula as

$$F_c/F_r > 1 - v/v'. \quad \text{(S7)}$$

Based on our experimental data in Figure 4, the furrow ingression rate of OptoMYPT-dark cells and OptoMYPT-pole cells were $v$ = 1.93 ± 0.33 µm/min and $v'$= 2.26 ± 0.25 µm/min, respectively. Thus, the cortical tension relative to ring tension is estimated as

$$F_c/F_r > 1 - 1.93/2.26 = 0.14. \quad \text{(S8).}$$

### Supplementary Figures

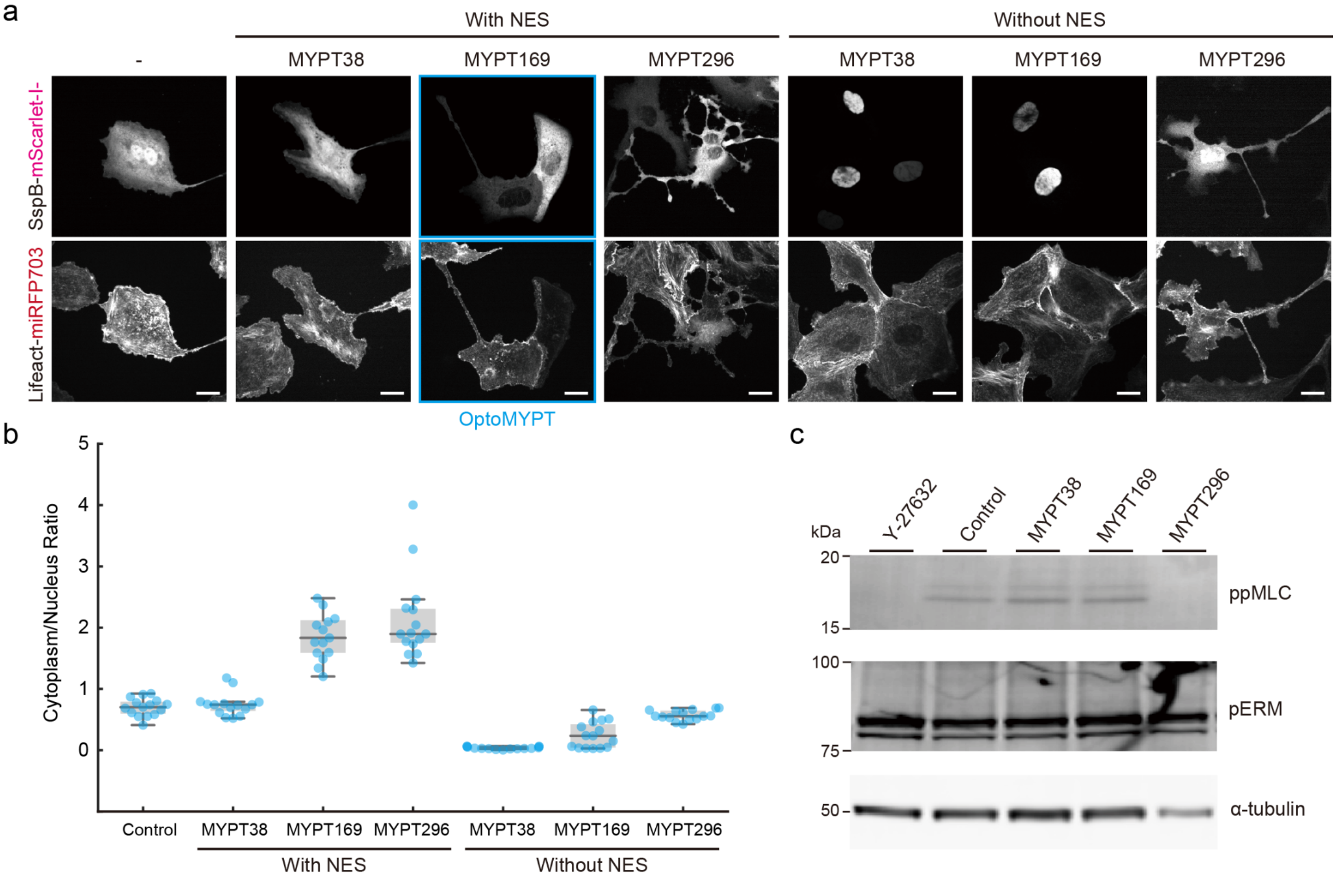


**Figure S1. Optimization of the length of PP1BD for OptoMYPT.**

(a) Representative images of MDCK cells transiently expressing SspB-mScarlet-I or SspB-mScarlet-I-PP1BDs with or without NES (upper panels), transiently expressing Stargazin-mEGFP-iLID, and stably expressing Lifeact-miRFP703 (lower panels). SspB-mScarlet-I-MYPT169-NES was used for the OptoMYPT (panels outlined in blue). Of note, SspB-mScarlet-I-MYPT296-expressing cells showed aberrant morphology with elongated protrusions. Scale bar, 20 µm. (b) The ratio of cytoplasmic to nuclear fluorescence intensity was quantified in each cell, and shown as a box plot. n = 14–15 cells. (c) Western blot analysis of ppMLC, pERM, and α-Tubulin in MDCK cells transiently expressing SspB-mScarlet-I (Control) or SspB-mScarlet-I-PP1BDs with NES, and Stargazin-mEGFP-iLID. Control cells were treated with 40 µM Y-27632 as a negative control (Y-27632).

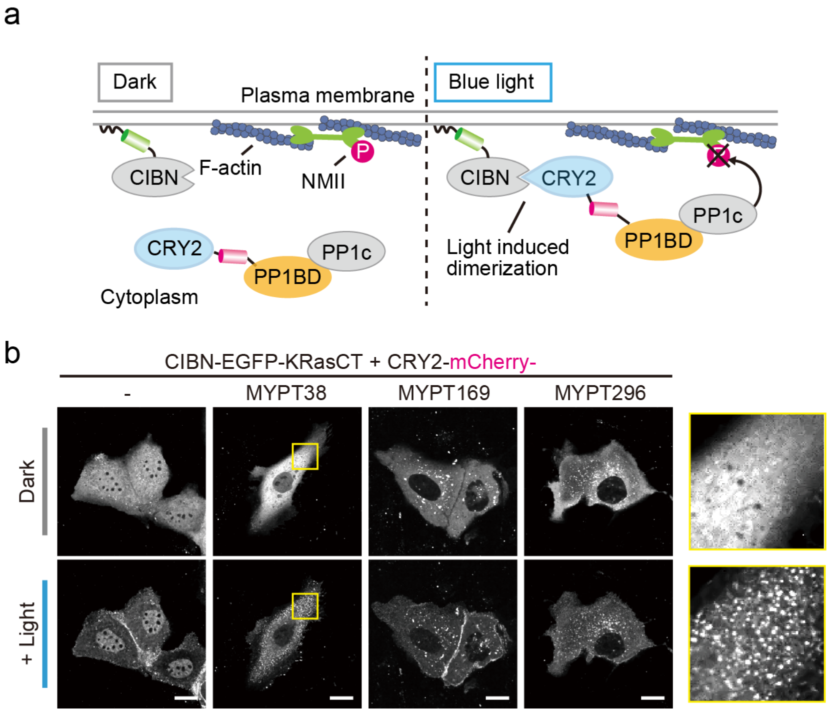


**Figure S2. CRY2-based OptoMYPT system.**

(a) Schematics of the CRY2-based OptoMYPT system. (b) Representative images of MDCK cells transiently expressing CRY2-mCherry or CRY2-mCherry-PP1BDs with NES, and CIBN-EGFP-KRasCT. The cells expressing CRY2-mCherry-MYPT38 showed aggregates and puncta upon blue light illumination (yellow boxed region). Scale bar, 20 μm.

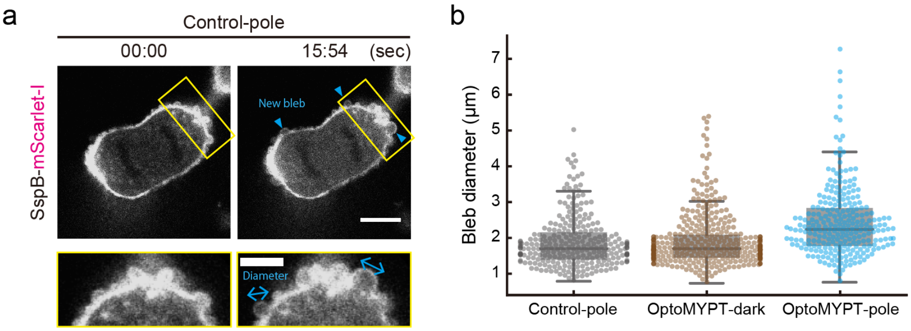


**Figure S3. Quantification of the bleb diameter.**

(a) Representative image of an MDCK cell expressing SspB-mScarlet-I and Stargazin-mEGFP-iLID during cytokinesis. The blue arrowheads in the upper panel indicate new blebs per 15.54 sec. The blue arrowheads in the lower panel indicate measured bleb diameters. Lower panels are the inset of yellow regions in upper panels. Scale bar, 10 and 5 µm for the upper and lower panels, respectively. (b) The diameter of blebs in the Control-pole, OptoMYPT-dark, and OptoMYPT-pole cells are shown as a box plot with swarm plot, in which each dot corresponds to a bleb diameter. The counted number of blebs was at least 250 for each sample.

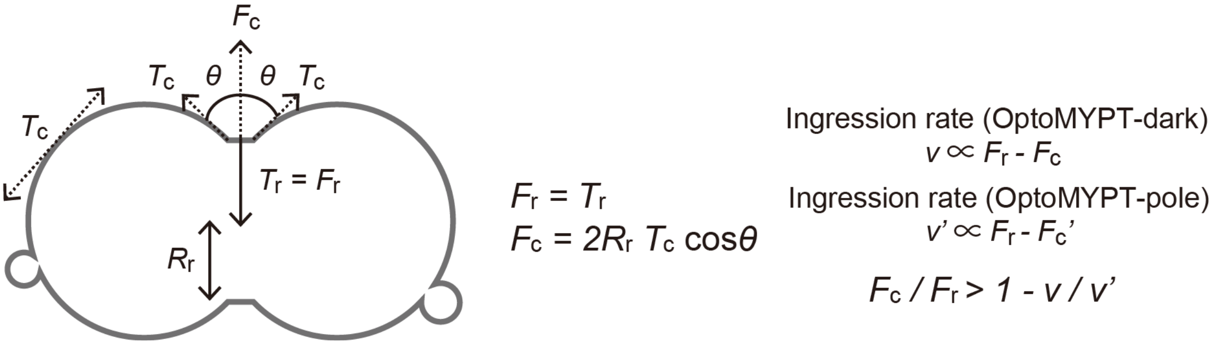


**Figure S4. Physical modeling.**

$T_c$ and $T_r$ are the cortical tensions at the actin cortex and contractile ring, respectively. $R_r$ is the ring diameter. $F_r$ and $F_c$ correspond to the force generated by ring tension and cortical tension, respectively.

#### Supplementary Movie legends

**Movie S1.** Membrane translocation of SspB-mScarlet-I-MYPT169 upon blue light illumination. Scale bar, 20 μm.

**Movie S2.** Membrane translocation of PP1c-miRFP703 upon blue light illumination. Scale bar, 20 μm.

**Movie S3.** Traction force measurement upon blue light illumination. Scale bar, 20 μm.

**Movie S4.** Induction of membrane protrusion by the OptoMYPT system. Blue rectangles indicate blue-light illuminated areas. Scale bar, 20 μm.

**Movie S5.** Induction of membrane retraction on the opposite side of the blue-light illuminated area. Scale bar, 20 μm.

**Movie S6.** Cytokinesis of a pole-illuminated MDCK cell expressing SspB-mScarlet-I. Scale bar, 10 μm.

**Movie S7.** Cytokinesis of a pole-illuminated MDCK cell expressing SspB-mScarlet-I-MYPT169. Scale bar, 10 μm.

**Movie S8.** Cytokinesis of an MDCK cell expressing SspB-mScarlet-I-MYPT169 under the dark condition. Scale bar, 10 μm.

#### Table S1. Plasmids used in this study.

| Plasmid name | Relevant features | Figures | Source or reference | Sequence |
|---|---|---|---|---|
| pCAGGS-SspB-mScarlet-I | CAG promoter, SspB-mScarlet-I | 1d-f, 3a,c,d,e, 4b-g, S1a,b, S3 | This study | https://benchling.com/s/seq-LizO8aJvYQvytA26tnvk |
| pCAGGS-SspB-mScarlet-I-MYPT38 | CAG promoter, SspB-mScarlet-I-MYPT38-NES | 1d-f, S1a,b | This study | https://benchling.com/s/seq-e5c4XNGJU7VUDY1Adw0D |
| pCAGGS-SspB-mScarlet-I-MYPT169 | CAG promoter, SspB-mScarlet-I-MYPT169-NES | 1d-f, 3b-e, 4b-g, S1a,b, S3 | This study | https://benchling.com/s/seq-VFtIByTEhOa69PaBJ7FF |
| pCAGGS-SspB-mScarlet-I-MYPT296 | CAG promoter, SspB-mScarlet-I-MYPT296-NES | 1d-f, S1a,b | This study | https://benchling.com/s/seq-NfHUqB2v72pMzw7Umbrv |
| pCAGGS-SspB-mScarlet-I-MYPT38 w/o NES | CAG promoter, SspB-mScarlet-I-MYPT38 (without NES) | S1a,b | This study | https://benchling.com/s/seq-M5vohEavn7YouFTOsbqR |
| pCAGGS-SspB-mScarlet-I-MYPT169 w/o NES | CAG promoter, SspB-mScarlet-I-MYPT169 (without NES) | S1a,b | This study | https://benchling.com/s/seq-4cIRYdnnkc5w1D7EG5pN |
| pCAGGS-SspB-mScarlet-I-MYPT296 w/o NES | CAG promoter, SspB-mScarlet-I-MYPT296 (without NES) | S1a,b | This study | https://benchling.com/s/seq-ik7oQqR9I1hNqtdzAjGK |
| pCAGGS-Stargazin-mEGFP-iLID | CAG promoter, Stargazin-mEGFP-iLID | 1d-f, 3a-e, 4b-g, S1a,b, S3 | This study | https://benchling.com/s/seq-XUhPNN7CavWnbGAq1TUe |
| pCAGGS-PP1c-miRFP703 | CAG promoter, PP1c-miRFP703 | 1d,f | This study | https://benchling.com/s/seq-8370yYN7igj1pyuf3TeZ |
| pCAGGS-CRY2-mCherry | CAG promoter, CRY2-mCherry | 2b,c, S2b | This study | https://benchling.com/s/seq-bhM5JFtyaGL3itlXyRYw |
| pCAGGS-CRY2-mCherry-MYPT38 | CAG promoter, CRY2-mCherry-MYPT38-NES | S2b | This study | https://benchling.com/s/seq-tLDYRgH228MpcEprZ8H3 |
| pCAGGS-CRY2-mCherry-MYPT169 | CAG promoter, CRY2-mCherry-MYPT169-NES | 2b,c, S2b | This study | https://benchling.com/s/seq-jlDVZxO6I2j5bRqXLNAe |
| pCAGGS-CRY2-mCherry-MYPT296 | CAG promoter, CRY2-mCherry-MYPT296-NES | S2b | This study | https://benchling.com/s/seq-KgPOub2LM388hRapgVNp |
| pCAGGS-CIBN-EGFP-KRasCT | CAG promoter, CIB N-terminus-EGFP-KRasCT | 2b,c, S2b | This study | https://benchling.com/s/seq-RA16GvytRbNXBNzKX1hg |
| pPBbsr2-Lifeact-miRFP703 | PiggyBac transposase donor vector, Lifeact-miRFP703 | 3a-e | This study | https://benchling.com/s/seq-6Umf325oqmk6HOmbykIX |
| pPBpuro-SspB-mCherry | PiggyBac transposase donor vector, SspB-mCherry | S1c | This study | https://benchling.com/s/seq-5d7Oj31YcZlxJqSsnNAa |
| pPBpuro-SspB-mCherry-MYPT38 | PiggyBac transposase donor vector, SspB-mCherry-MYPT38-NES | S1c | This study | https://benchling.com/s/seq-fZysCS2LhI69PbhGD4tg |

| | | | | |
|---|---|---|---|---|
| pPBpuro-SspB-mCherry-MYPT169 | PiggyBac transposase donor vector, SspB-mCherry-MYPT169-NES | S1c | This study | https://benchling.com/s/seq-RrmRNQx2r9ULkAGO3yxd |
| pPBpuro-SspB-mCherry-MYPT296 | PiggyBac transposase donor vector, SspB-mCherry-MYPT296-NES | S1c | This study | https://benchling.com/s/seq-WFNiQng3AysDppDUz4fk |
| pPBbsr2-Stargazin-mEGFP-iLID | PiggyBac transposase donor vector, Stagazin-mEGFP-iLID | S1c | This study | https://benchling.com/s/seq-iKCLTeMJ2f84JqYhfn1n |
| pCAGGS-hyPBase | CAG promoter, PiggyBac transposase | 3a-e, S1c | This study | https://benchling.com/s/seq-oGkw53b41IZqvzF5yQ9K |
